# An Integrated Analysis of Network Pharmacology, Molecular Docking, and Experiment Validation to Explore the Mechanism of Danbai Granules in Treating Sequelae of Pelvic Inflammatory Disease

**DOI:** 10.1101/2023.03.05.531232

**Authors:** Yuying Zhu, Xiaoou Xue, Jun Li, Zilin Long, Jiawei Zhang, Baohua Zhao

## Abstract

**Purpose:** To explore the possible pharmacological mechanisms of Danbai Granules (DBG) against sequelae of pelvic inflammatory disease (SPID) by the method of network pharmacology and molecular docking combined with experimental validation.

**Methods:** Firstly, biologically active compounds, targets of DBG, and related targets of SPID were collected from public databases. The protein– protein interaction (PPI) network, Gene GO and KEGG enrichment analysis were carried out via R language to find the potential biological processes and pathways of DBG against SPID. Subsequently, molecular docking was used to calculate the affinity between the top six targets and the top ten compounds, respectively. Finally, the above analysis results were verified by in vivo experiments.

**Results:** A total of 106 crossover genes were selected from 220 potential medicine genes and 952 SPID-related genes. PPI network analysis indicated that IL-6, IL-1β, VEGFA, CASP3, PTGS2, JUN, etc., were core targets. GO and KEGG enrichment analysis revealed that PI3K-Akt, AGE–RAGE, IL-17, TNF, HIF-1 and Toll-like receptors signaling pathways were the main signaling pathways. Molecular docking showed that 15 pairs of target-ingredient combinations have good binding activity. The animal experiments showed that DBG administration inhibited the expression of TLR4/MyD88/NF-κB p65 signaling pathway-related proteins expression, altered the high expression of IL-6, IL-1β, and low level of IL-4, IL-10, thereby relieving the pelvic inflammation of SPID.

**Conclusion:** Through the network pharmacology and molecular docking, our findings predicted the active ingredients and potential targets of DBG in treating SPID. The experimental verification preliminarily revealed that DBG may inhibit the inflammatory response of SPID rats by regulating the TLR4/MyD88/NF-κB p65 signaling pathway. which provides a theoretical basis for the study of the pharmacodynamic material basis and mechanism of DBG against SPID at the comprehensive level.

## Introduction

Pelvic inflammatory disease (PID) refers to a group of infectious diseases of the female upper genital tract, mainly including endometritis, salpingitis, tubo-ovarian abscess, and pelvic peritonitis. If pelvic inflammatory disease is not properly diagnosed or treated in time, sequelae of pelvic inflammatory disease (SPID) may occur. and its main pathological changes are tissue destruction, extensive adhesion, hyperplasia and scarring, which may lead to infertility, ectopic pregnancy, chronic pelvic pain, recurrent pelvic inflammatory disease, etc [1–3]. The disease often occurs repeatedly, and it is difficult to heal. The incidence rate is increasing year by year, and there is a trend of younger people. The literature shows that the incidence of pelvic inflammatory disease can reach 2% to 12% [4–5], which seriously affects the life quality and reproductive health of women. At present, modern medicine mainly adopts antibiotic therapy, analgesic therapy, physical therapy, surgery, immunotherapy, etc. based on clinical manifestations, pathogens and various pathogenesis hypotheses. However, these treatment methods have drawbacks such as antibiotic resistance, adverse effects, and disease recurrence. Therefore, formulating more effective SPID treatment strategies to reduce the occurrence of side effects and prevent disease recurrence has an important clinical guiding role.

Proven by thousands of years of clinical practice and supported by research evidence on the superiority of TCM clinical RCTs, TCM can achieve multi-channel comprehensive intervention for SPID and achieve definite clinical efficacy under the guidance of the “holistic view” theory. Therefore, traditional Chinese medicine therapy has unique advantages and broad clinical application prospects in the prevention and treatment of SPID. Danbai Granules is one of the commonly used Chinese patent medicines for the treatment of SPID, which contains 13 kinds of Chinese medicines: Moutan Cortex(MDP), Sargentodoxae Caulis (DXT), Violae Herba (ZHDD), Sparganii Rhizoma (SL), Curcumae Rhizoma (EZ), Herba Patrinae (BJC), Chuanxiong Rhizoma (CX), Paeoniae Radix Alba (BS), Smilacis Glabrae Rhizoma (TFL), Herba Solani Lyrati (BY), Herba Hedyotidis (BHSSC), Patrinia Heterophylla Bunge (MTH), Ailanthi Cortex (CP), Angelica Sinensis (DG) . It has the functions of clearing heat and removing blood stasis, removing dampness and relieving pain, which is in line with the fundamental pathogenesis of damp-heat stasis of sequelae of pelvic inflammatory disease. Long-term clinical observation has proven its effect in relieving pelvic pain and treating SPID, However, the mechanism of DBG for treatment of SPID remains inconclusive. Therefore, further research is required to explore the underlying mechanisms involved.

Traditional Chinese medicine and its compounds have the unique functions of multi-ingredient, multi-channel, multi-target and multi-effect. And due to its integral therapeutic effects, it makes the study of the material basis, in vivo processes, biological effects and their interrelationships of traditional Chinese medicines and their compounds quite complicated, and it is a challenge to figure out its pharmacological and molecular mechanisms on diseases. Therefore, it is very necessary to explore new strategies and methods to study the mechanism of action of traditional Chinese medicine and its compounds [6]. In recent years, network pharmacology has been used more and more widely in the research of traditional Chinese medicine prescriptions, and it is recognized as one of the important methods of researching traditional Chinese medicine prescriptions. Network pharmacology is a new subject that integrates systems biology pharmacology and bioinformatics to explore the complex mechanisms of drugs from a multidimensional perspective using a “disease-target-herb” network model [7–9]. Hence, network pharmacology based on large databases has become a powerful tool for TCM research. Luanqian reveals the anti-inflammatory mechanism exhibited by Radix chinensis and Radix Paeoniae Alba in SPID through network pharmacology and experimental validation [10]. In addition, molecular docking can simulate molecular geometry and intermolecular forces by stoichiometric methods [11]. At present, many researchers have applied network pharmacology and molecular docking methods to the study of traditional Chinese medicine in the treatment of various diseases, and have achieved corresponding results, enriching the connotation of traditional Chinese medicine [12–13].

In the present study, using the method of network pharmacology, we obtained the pharmacodynamic substances of DBG and the co-acting targets of DBG and SPID. Subsequently, we constructed a multi-dimensional network map, PPI network, GO functional enrichment and KEGG signaling pathway, and combined molecular docking to investigate the complex mechanism of action of DBG on SPID. Finally, the predicted major target proteins and Toll-like receptor signaling pathway were further verified in vivo experiments. These findings can provide theoretical basis and scientific support for follow-up research. A flowchart of this research is shown in Fig 1.

**Fig 1.**
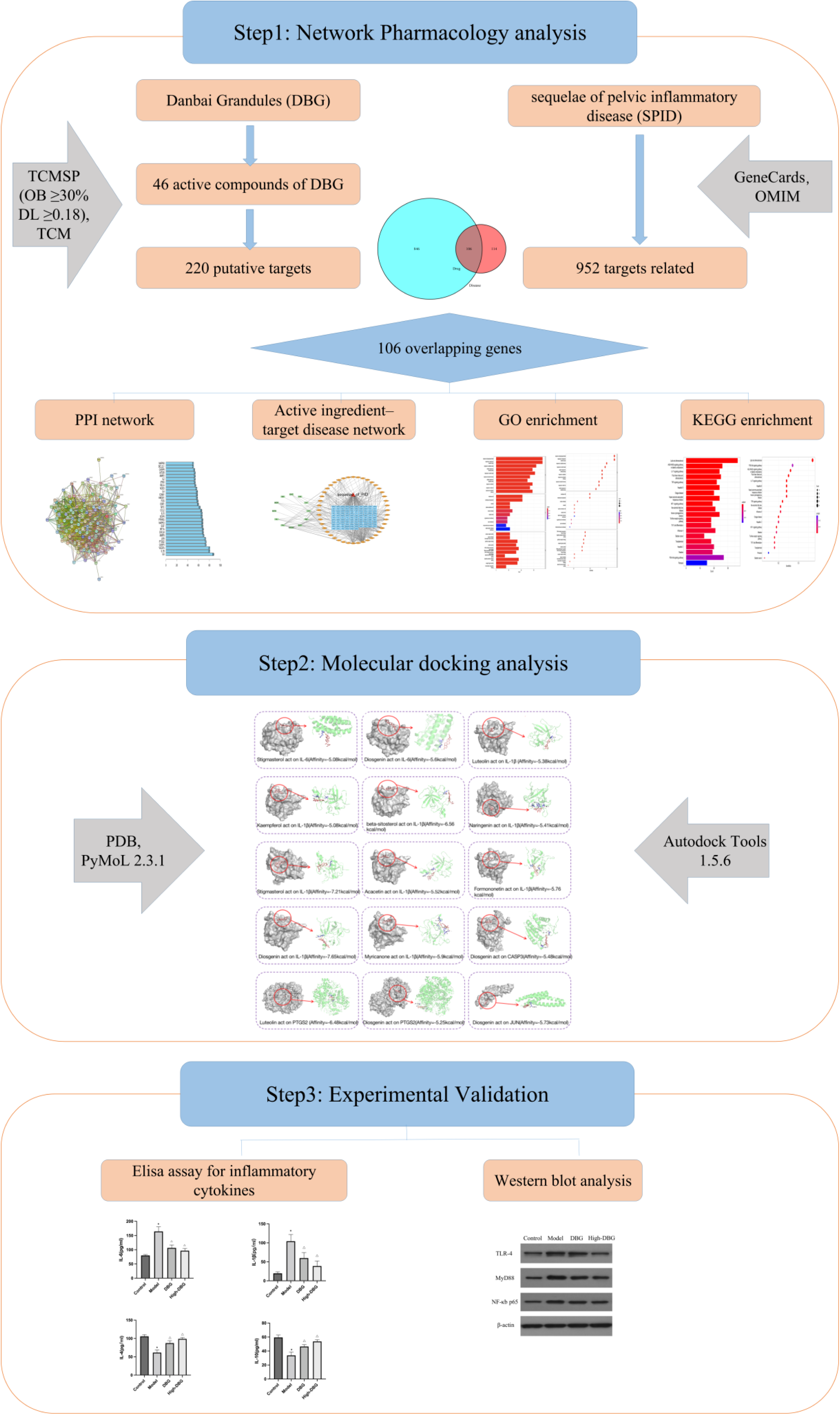
The detailed flowchart of the study design

## Materials and Methods

### Main Component Screening

The main chemical compounds of the DBG were collected through the Traditional Chinese Medicine Systems Pharmacology database (TCMSP, https://old.tcmsp-e.com/tcmsp.php) [14], According to the pharmacokinetic parameters of the compounds, the criteria of oral bioavailability (OB) ≥ 30% and drug-like properties (DL) ≥ 0.18 were set to screen out candidate active ingredients [15]. Targets corresponding to the candidate compounds of DBG were retrieved through TCMSP and TCM databases (TCM, http://tcm.cmu.edu.tw) [16] and combined with Drug Bank (https://www.drugbank.ca). [17] Subsequently, query the gene name corresponding to the target through the Uniprot database (https://www.uniprot.org/) [18], select the species as “Homo sapiens” to establish a dataset, and the target protein names corresponding to the Uniprot dataset were uniformly converted into gene names. Then, search the GeneCards online database (https://www.genecards.org/) [19] and OMIM (http://omim.org/) [20] using the keyword “Sequelae of Pelvic Inflammatory Disease” and “Chronic Pelvic Inflammatory Disease” to obtain a dataset of SPID-related targets. Finally, we constructed a Venn diagram to display DBG and SPID intersection genes using R software 4.1.2. These may be potential targets of DBG for the treatment of SPID.

### Construction and analysis of the protein– protein interaction network

We inputted these overlapping targets into STRING (https://string-db.org) [21] database to construct the PPI network. The species was set to “Homo sapiens”, the minimum interaction threshold was “medium confidence (0.400)”, a single isolated node was eliminated, and other parameters remained unchanged to obtain key target interaction information. The obtained protein interaction results were saved in TSV format. The count.R plugin in the R language software was applied to find out the core targets.[22] The top 20 closely related genes were screened.

### Network construction of drugs-compounds-genes-disease

The interacting genes were obtained from target genes extracted from DBG active components and those of SPID. Then, we used the Cytoscape software 3.9.0 (https://cytoscape.org/) [23] for visual analysis of these data to draw the interaction network. The medicinal materials, ingredients, and targets are represented by nodes in the network graph, and the interaction between two nodes is represented by edges. The Network Analyzer tool [24] in Cytoscape software was used for calculating the degree value of node.

### Gene Ontology and Kyoto Encyclopedia of Genes and Genomes Pathways Enrichment Analysis

To obtain the biological process, cellular components, molecular functions and related signaling pathways of DBG in the treatment of SPID. we performed GO and KEGG enrichment analysis by using the cluster Profiler Go.R plug-in in R software [25] and Perl script [26]. Our requirements for setting the visualization results are “width=35, height=20, units=“cm”. The p-value and q-value were set to less than 0.05. Then, the cross-targets were analyzed by biological processes (BP), cellular component (CC), molecular function (MF) and KEGG pathway, and the bar graph maps were drawn to predict the specific mechanism of the DBG against SPID.

### Molecular Docking of active ingredients and core targets

To verify the binding activity between the components and core targets of screened by network pharmacology, molecular docking was performed for the top ten ingredients and the top six targets. First, the three-dimensional structure of the main components of DBG was searched in the TCMSP database as a small molecule ligand file for molecular docking. Subsequently, the protein ID (species human) corresponding to the screened network core genes was searched in the UniProt database, and the protein ID was imported into the PDB database (https://www.rcsb.org/) [27], protein database to obtain the three-dimensional structure of the protein, which was used as the macromolecular receptor file for molecular docking. Then, the PyMOL (2.5.1) [28] software was used to remove the water molecules and small molecule ligands of the protein, and the ligands and receptors were respectively hydrogenated with the help of AutoDock Vina (1.1.2) software for docking [29]. Only the free energy (kcal·mol-1) of each pair of molecular docking was retained, and the docking results were finally visualized using PyMOL (2.5.1) software. It is generally believed that the binding energy is ≤ 0, indicating that small molecule ligands can spontaneously bind to protein receptors. When the binding energy is ≤ -5.0 kcal/mol, it is considered to have strong binding activity, which also suggests that the key targets predicted by network pharmacology had high reliability [30].

## Experimental Validation

### Animals

A total of 36 8-week-old female rats (weighing 200± 20g) were obtained from Beijing Weitong lihua Co., Ltd. (Beijing, China, SCXK(Beijing) : 2016-0006). The rats were raised in the SPF animal laboratory, including room temperature of 18∼25°C, humidity at 60%∼80%, and a 12h daily light exposure. They were adaptively fed for 1 week. The experimental protocols and ethics were approved by the Institutional Animal Care and Use Committee of Dongzhimen Hospital affiliated to Beijing University of Chinese Medicine (No. BUCMDZM-2021-40). and all laboratory procedures were performed in accordance with the Regulations for the Regulations on the Administration of Laboratory Animals approved by the State Council of the People’s Republic of China.

### Drug and Reagents

Danbai Granule was acquired from Shandong Lunan Houpu Co., Ltd.(Zhejiang, China). It contains Moutan Cortex, Sargentodoxae Caulis, Violae Herba, Sparganii Rhizoma, Curcumae Rhizoma, Herba Patrinae, Chuanxiong Rhizoma, Paeoniae Radix Alba, Smilacis Glabrae Rhizoma, Herba Solani Lyrati, Herba Hedyotidis, Patrinia heterophylla Bunge, Ailanthi Cortex, and Angelica Sinensis.

Rats IL-6, IL-1 β, IL-4, IL-10 ELISA Kits were purchased from Shanghai Enzyme Link Biotechnology Co., Ltd.(Shanghai, China). Rabbit TLR-4 Antibody, MyD88 Antibody and NF -kB p65 Antibody were obtained from Abcam Company Ltd. (Cambridge, MA, USA).

### SPID Model

The rats model of SPID was established according to previous experiments [31] by mechanical damage combined with 25% phenol mortar injection at a dose of 0.06ml. The rats exhibited the characteristics of SPID in terms of uterus pathology [32].

### Animal Grouping, Treatment, and Specimen collection

36 female rats were randomly divided into 4 groups with 9 rats in each group: Model Group, Control Group, DBG Group (2.5g/kg/d), High-dose DBG Group (5g/kg/d). Model Group, DBG groups were first intraperitoneally administered with 25% phenol mortar injection at a dose of 0.06ml to induce SPID, After successful modeling, each DBG group was administered the corresponding dose of drugs respectively, Meanwhile, the model group and the control group were given normal saline, once a day, for 2 consecutive weeks. Rats in all groups were intragastrically administered at a dose of 10 mL/kg. The dosage of the drug in rats is converted according to the body surface area conversion algorithm [33]. The rats were fasted on the morning of the 2nd day after the last administration. Then, they were anesthetized and 4-5 mL of blood was collected. Subsequently, uterine tissue was collected after the rats were sacrificed. The animal experiments were approved by The Animal Ethics Committee of Beijing University of Traditional Chinese Medicine (Beijing, China).

### Elisa assay for Inflammatory Cytokines in Each Group

After the treatment, blood was drawn from rat abdominal aorta and serum was obtained by centrifugation at 3500 rpm for 10 min. IL-6, IL-1β, IL-4 and IL-10 levels in serum were measured using ELISA kit according to the manufacturer’s protocol.

### Western Blot Analysis

The influence of DBG on the expressions of TLR-4, MyD88, NF-κb p65 in protein level was assessed by Western blot. Taking 0.05g of cryopreserved uterine tissues in centrifuge tubes, total protein of which was routinely extracted with tissue lysate, and protein concentration was determined with BCA protein concentration determination kit. The protein was normalized to 5 mg/mL and loaded onto SDS-PAGE gels for electrophoresis. After electrophoresis, the protein bands isolated from the gel were transferred to the PVDF membrane by transfer electrophoresis. 5% skimmed milk powder solution was used for blocking 1 h at room temperature, washed with TBST, and TLR4, MyD88, NF-κB p65, and β-actin primary antibodies (dilution ratio 1:1000) was added and incubated overnight in a refrigerator at 4 °C. After washing again with TBST solution, universal secondary antibody was added and then incubated at room temperature for 1.5 h. Membranes were washed with TBST solution and developed with ECL luminescent solution. The film was exposed in a dark room with X-ray film, developed and fixed, then the film was scanned, and the optical density of the target band was analyzed using Quantity One image analysis software.

### Statistical Analysis

Statistical analysis was performed using SPSS 26.0 software, The data are expressed as mean ± SD. One-way ANOVA was used for the comparison of means among the groups [34], P < 0.05 was considered statistically significant. When p<0.05, it was considered to be statistically significant, and the difference between groups was significant.

## Results

### Targets of DBG against SPID

Excluding no known targets of compounds, A total of 46 ingredients were identified. Of these ingredients, 6 were from Moutan Cortex, 4 from Sargentodoxae Caulis, 5 from Sparganii Rhizoma, 1 from Curcumae Rhizoma, 11 from Herba Patrinae, 6 from Chuanxiong Rhizoma, 8 from Paeoniae Radix Alba, 14 from Smilacis Glabrae Rhizoma, 5 from Herba Hedyotidis, 1 from Patrinia Heterophylla Bunge, 2 from Angelica Sinensis, and 11 from Ailanthi Cortex. The selected 46 compounds are shown in Table 1. After searching and removing duplicate targets using the TCMSP database, TCM and Drug Bank, all active compounds had 220 targets. Then, we use UniProt database and selected species option as “Homo sapiens” to acquire all targets after removing the repetitive targets and we converted the target protein name to a gene name. Moreover, a total of 952 SPID-related targets were identified from the Gene Cards database. Ultimately, a total of 106 intersecting genes, also known as potential genes, were gathered for the purpose of studying the various processes underlying the action of DBG in the treatment of SPID are shown in Fig 2.

**Fig 2.**
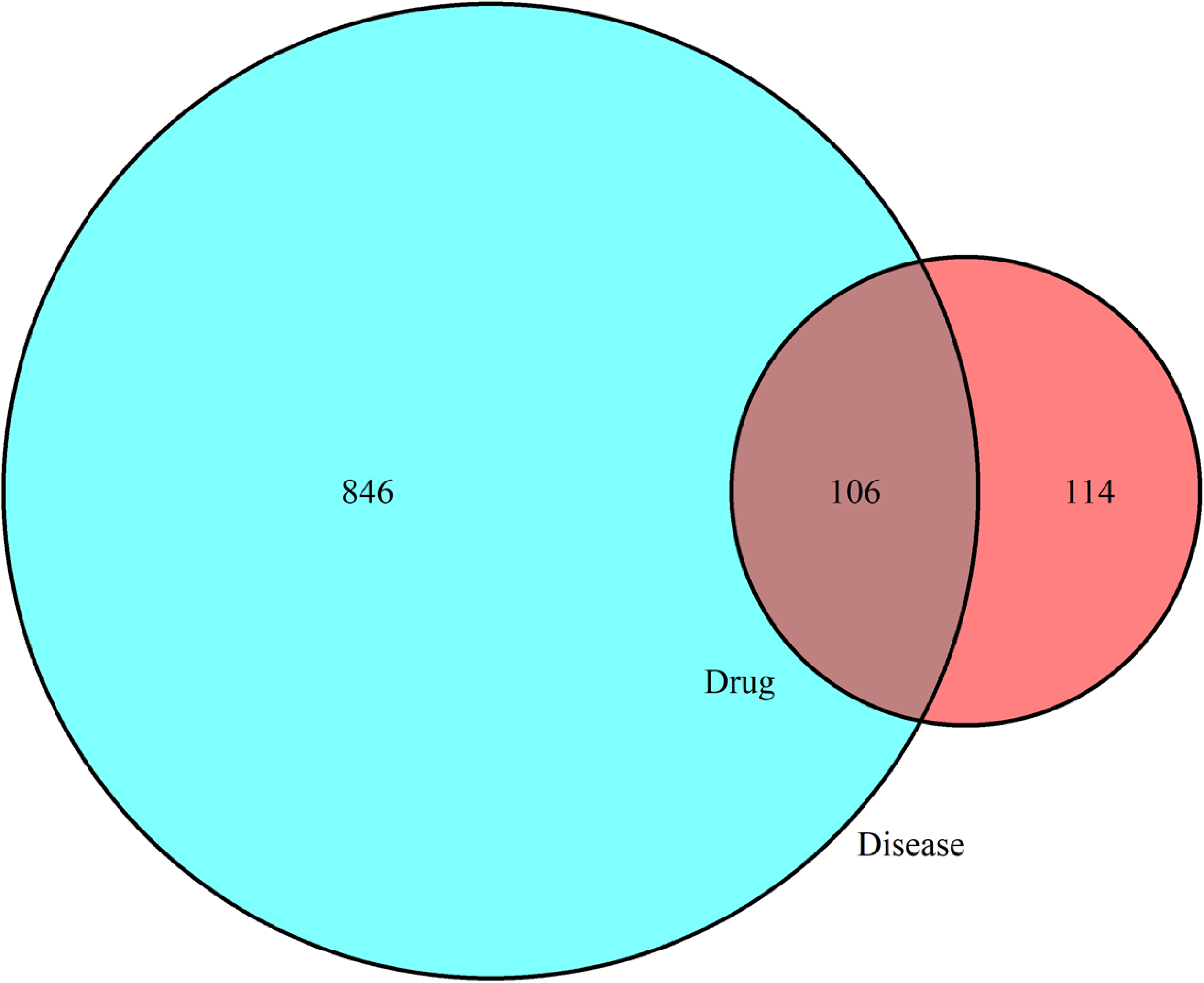
Venn diagram of the DBG (drug genes) and pelvic inflammatory disease (disease genes) intersection targets

**Table 1:** Molecular ID, molecule name, oral availability, AND drug-Like Properties Of 46 ingredients of DBG

| Molecular ID | Molecular name | Oral | Dru | Herb |
| --- | --- | --- | --- | --- |

|  |  | Availa-bility | g-Li<br>ke |  |
| --- | --- | --- | --- | --- |
| MOL000211 | Mairin | 55.38 | 0.78 | MDP, BS |
| MOL000359 | Sitosterol | 36.91 | 0.75 | MDP, BJC,<br>CX, BS, TFL |
| MOL000422 | Kaempferol | 41.88 | 0.24 | MDP, BJC,<br>BS, CP |
| MOL000492 | (+)-catechin ( ( + ) ) | 54.83 | 0.24 | MDP, BS |
| MOL007374 | 5-[[5-(4-methoxyphenyl)-2-furyl]methyle<br>ne]barbituric acid | 43.44 | 0.3 | MDP |
| MOL000098 | quercetin | 46.43 | 0.28 | MDP, BJC,<br>TFL, BHSSC,<br>MTH, CP |
| MOL007923 | 2-(4-hydroxyphenyl)<br>ethyl<br>(E)-3-(4-hydroxyphenyl)prop-2-enoate | 93.36 | 0.21 | DXT |
| MOL000096 | (-)-catechin ( - ) | 49.68 | 0.24 | DXT |
| MOL000358 | beta-sitosterol | 36.91 | 0.75 | DXT, SL,<br>BJC, BS,<br>TFL,<br>BHSSC, DG,<br>CP |
| MOL001297 | trans-gondoic acid | 30.7 | 0.2 | SL |
| MOL000296 | Hederagenin | 36.91 | 0.75 | SL |
| MOL000392 | Formononetin | 69.67 | 0.21 | SL |
| MOL000449 | Stigmasterol | 43.83 | 0.76 | SL, BJC, TFL,<br>BHSSC, DG,<br>CP |
| MOL001689 | Acacetin | 34.97 | 0.24 | BJC |
| MOL001677 | asperglaucide | 58.02 | 0.52 | BJC |
| MOL001678 | bolusanthol B | 39.94 | 0.41 | BJC |
| MOL002322 | Isovitexin | 31.29 | 0.72 | BJC |
| MOL000006 | Luteolin | 36.16 | 0.25 | BJC |
| MOL001697 | Sinoacutine | 63.39 | 0.53 | BJC |
| MOL001494 | Mandenol | 42.00 | 0.19 | CX |
| MOL002135 | Myricanone | 40.60 | 0.51 | CX |
| MOL002140 | Perlolyrine | 65.95 | 0.27 | CX |
| MOL002157 | wallichilide | 42.31 | 0.71 | CX |
| MOL000433 | FA | 68.96 | 0.71 | CX |
| MOL001918 | paeoniflorgenone | 87.59 | 0.37 | BS |
| MOL001919 | (3S,5R,8R,9R,10S,14S)-3,17-dihydroxy-4,4,8,10,14-pentamethyl-2,3,5,6,7,9-hexahydro-1H-cyclopenta[a]phenanthrene-15,16-dione | 43.56 | 0.53 | BS |
| MOL001924 | paeoniflorin | 53.87 | 0.79 | BS |
| MOL013117 | 4,7-Dihydroxy-5-methoxyl-6-methyl-8-formyl-flavan | 37.03 | 0.28 | TFL |
| MOL013119 | Enhydrin | 40.56 | 0.74 | TFL |
| MOL013129 | (2R,3R)-2-(3,5-dihydroxyphenyl)-3,5,7-t | 63.17 | 0.27 | TFL |
|  | rihydroxychroman-4<br>-one |  |  |  |
| MOL001736 | (-)-taxifolin | 60.51 | 0.27 | TFL |
| MOL004328 | naringenin | 59.29 | 0.21 | TFL |
| MOL004567 | isoengelitin | 34.65 | 0.70 | TFL |
| MOL004575 | astilbin | 36.46 | 0.74 | TFL |
| MOL004576 | taxifolin | 57.84 | 0.27 | TFL |
| MOL004580 | cis-Dihydroquercetin | 66.44 | 0.27 | TFL |
| MOL000546 | Diosgenin | 80.88 | 0.81 | TFL |
| MOL001659 | Poriferasterol | 43.83 | 0.76 | BHSSC |
| MOL001670 | 2-methoxy-3-methyl<br>-9,10-anthraquinone | 37.83 | 0.21 | BHSSC |
| MOL001507 | stigmast-4-en-3-one | 36.08 | 0.76 | CP |
| MOL001755 | 24-Ethylcholest-4-en<br>-3-one | 36.08 | 0.76 | CP |
| MOL006277 | Shinjudilactone | 43.56 | 0.74 | CP |
| MOL006279 | Shinjulactone B | 98.40 | 0.47 | CP |
| MOL006303 | 4,5-dihydrocanthin-6 | 48.58 | 0.28 | CP |
|  | -one |  |  |  |
| MOL006304 | 4-hydroxy-canthin-6<br>-one | 35.11 | 0.25 | CP |
| MOL006311 | Artelin | 41.23 | 0.22 | CP |

### Results of Network

#### Results of PPI Network

We used the STRING online database service platform to construct the PPI network. This PPI network consists of 106 nodes and 1968 edges, with an average node degree of 37.1 and an average local clustering coefficient of 0.675 (as shown in Fig 3A).Then, R language software was used to calculate the top 30 targets of frequency and draw a bar graph (as shown in Fig 3B), in which IL-6, IL-1β, VEGFA, CASP3, PTGS2 and JUN are the whole core proteins, indicating that Danbai granules may play a therapeutic role through these proteins in the treatment of sequelae of pelvic inflammatory disease.

**Fig 3.**
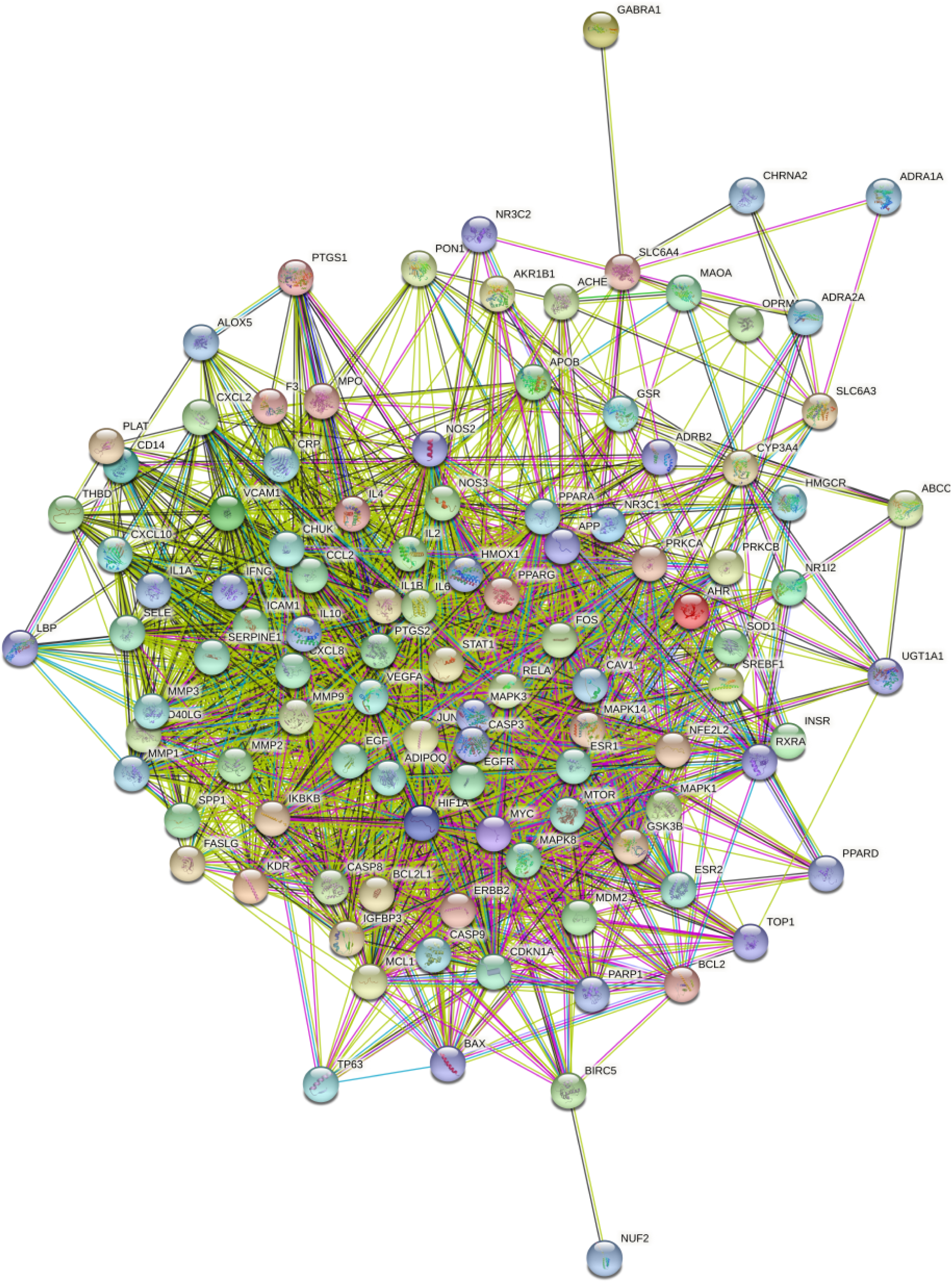
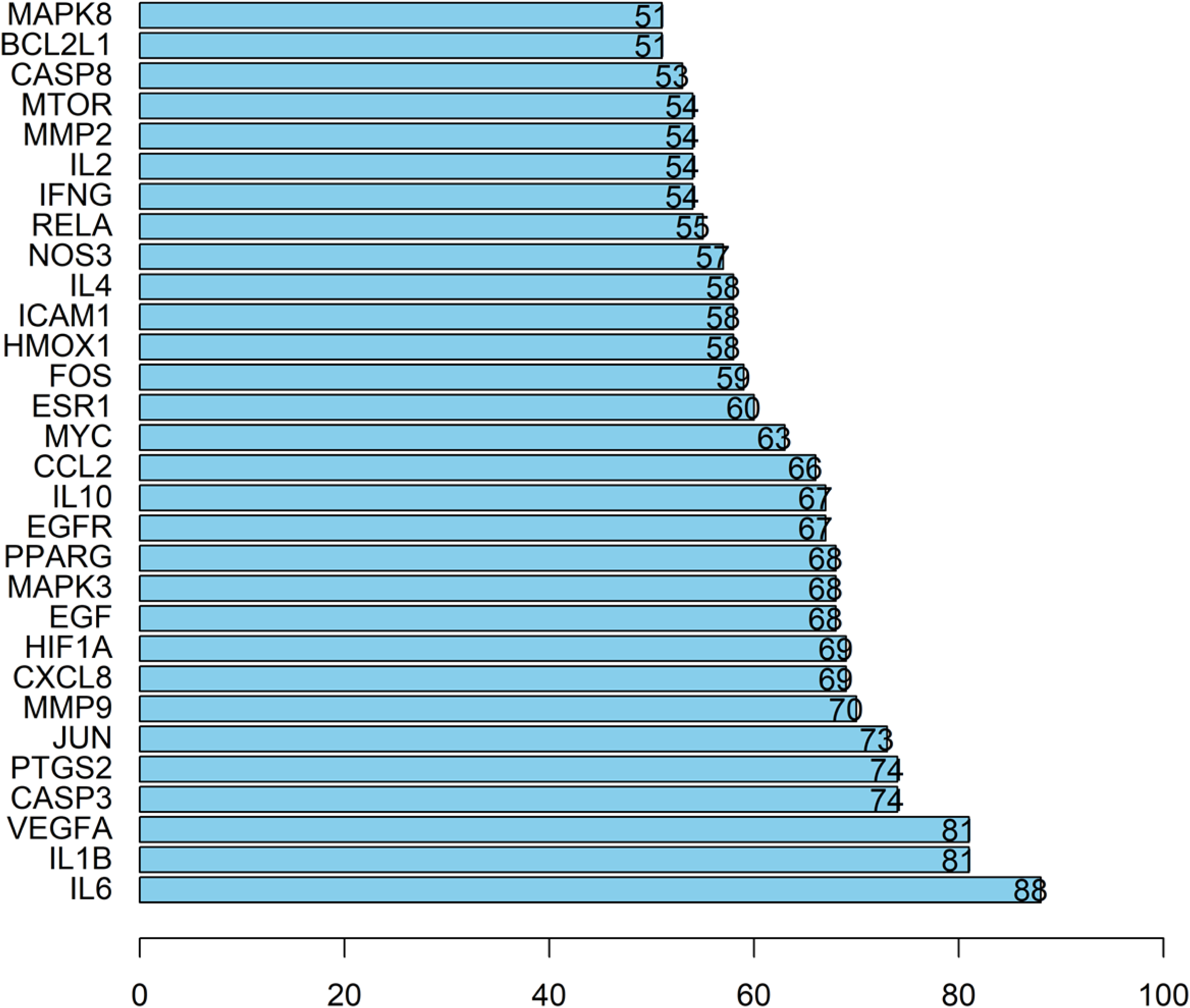
A. Protein interaction network of DBG in the treatment of SPID B. Frequency display of PPI in the treatment of SPID with DBG (x-axis the number of edges joined by the target)

Cytoscape Pharmacological Network Diagram of Targets and the Result of Hub Gene Selection Cytoscape 3.9.0 and Perl language were used to construct a four-dimensional regulatory network of DBG-drug substance-target-SPID, with 150 nodes and 500 edges (as shown in Fig 4). Then, we set the node color and shape to show the drug-disease interaction. The red triangle represents SPID, green rectangles represent drugs of DBG, blue rectangles represent the common targets, and orange ovals represent molecular ID.

**Fig 4.**
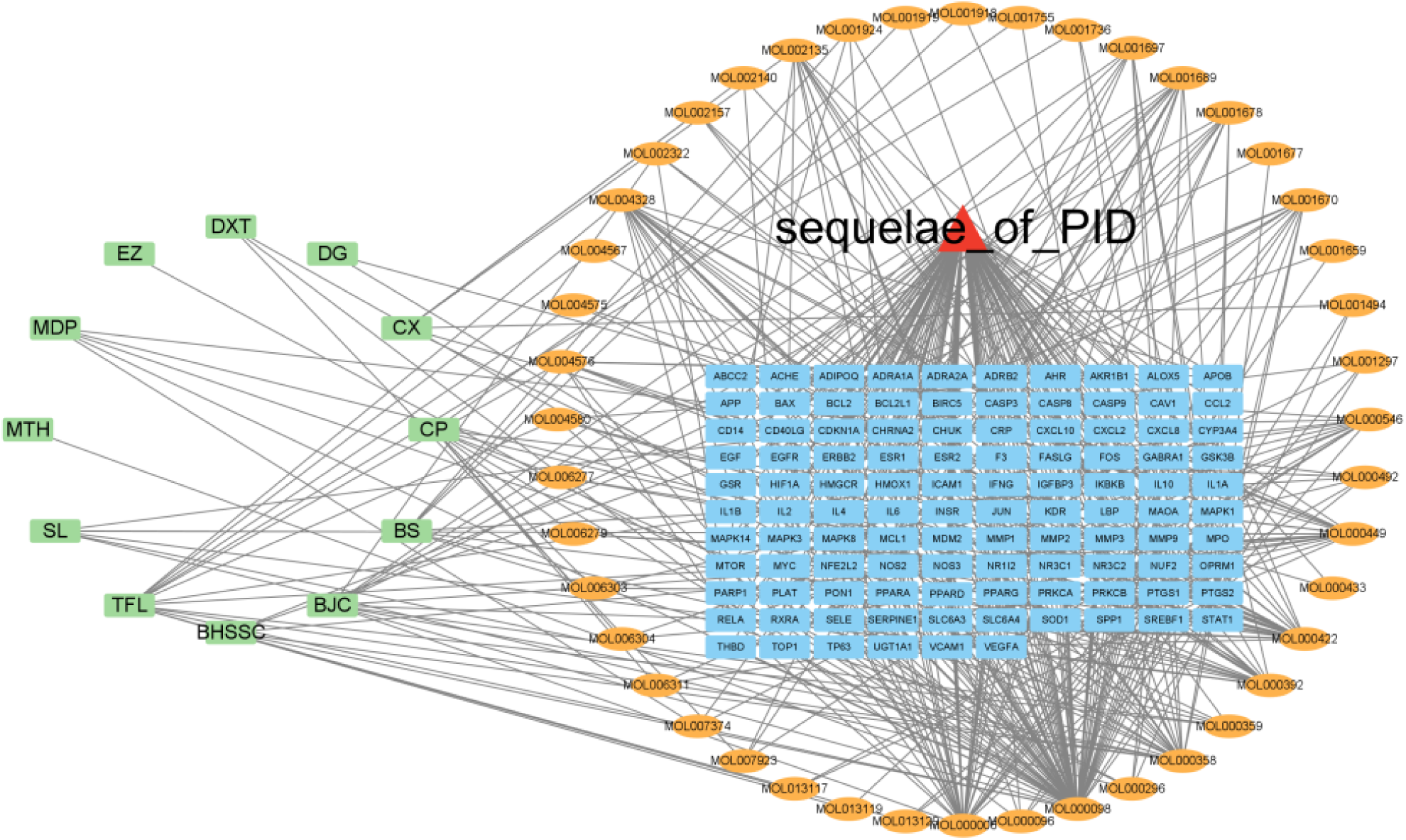
Active ingredient–target disease network of DBG in the treatment of SPID

Network topology analysis is performed based on topological data such as degree, betweenness, and closeness associated with the “Network Analyzer” section. According to the degree of each node in the network graph, the main active components of DBG used to control SPID are screened. The top ten active ingredients are MOL000098 (quercetin), MOL000006 (Luteolin), MOL000422 (Kaempferol), MOL000358 (beta-sitosterol), MOL004328 MOL000422 (Kaempferol), MOL000358 (beta-sitosterol), MOL004328 (naringenin), MOL000449 (stigmasterol), MOL001689 (acacetin), MOL000392 (formononeti), MOL000546 (diosgenin), MOL002135 (Myricanone).

### Results of Enrichment Analysis

#### GO Analysis

Using Cluster Profiler GO.R plug-in and Perl script to perform GO functional enrichment analysis on the 106 common target genes screened, we got 2978 GO entries, of which 2593 GO entries were related to biological process (BP). The enrichment results indicate that the mechanism of DBG against SPID may be related to the response to lipopolysaccharide, molecule of bacterial origin, oxidative stress, chemical stress, nutrient levels. The cell component (CC) pathway was enriched to obtain 170 related GO items, which had significant effects on cellular components such as membrane, caveloa, nuclear envelope and vesicle lumen, Moreover, the molecular function (MF) pathway was enriched to obtain 215 items, which mainly included DNA-binding transcription factor banding, signaling receptor activator activity, cytokine activity, receptor ligand activity. Among the above entries, the top 10 entries with P<0.05 in BP, CC, and MF were screened and visualized, as shown in Figs 5A and 5B.

**Fig 5.**
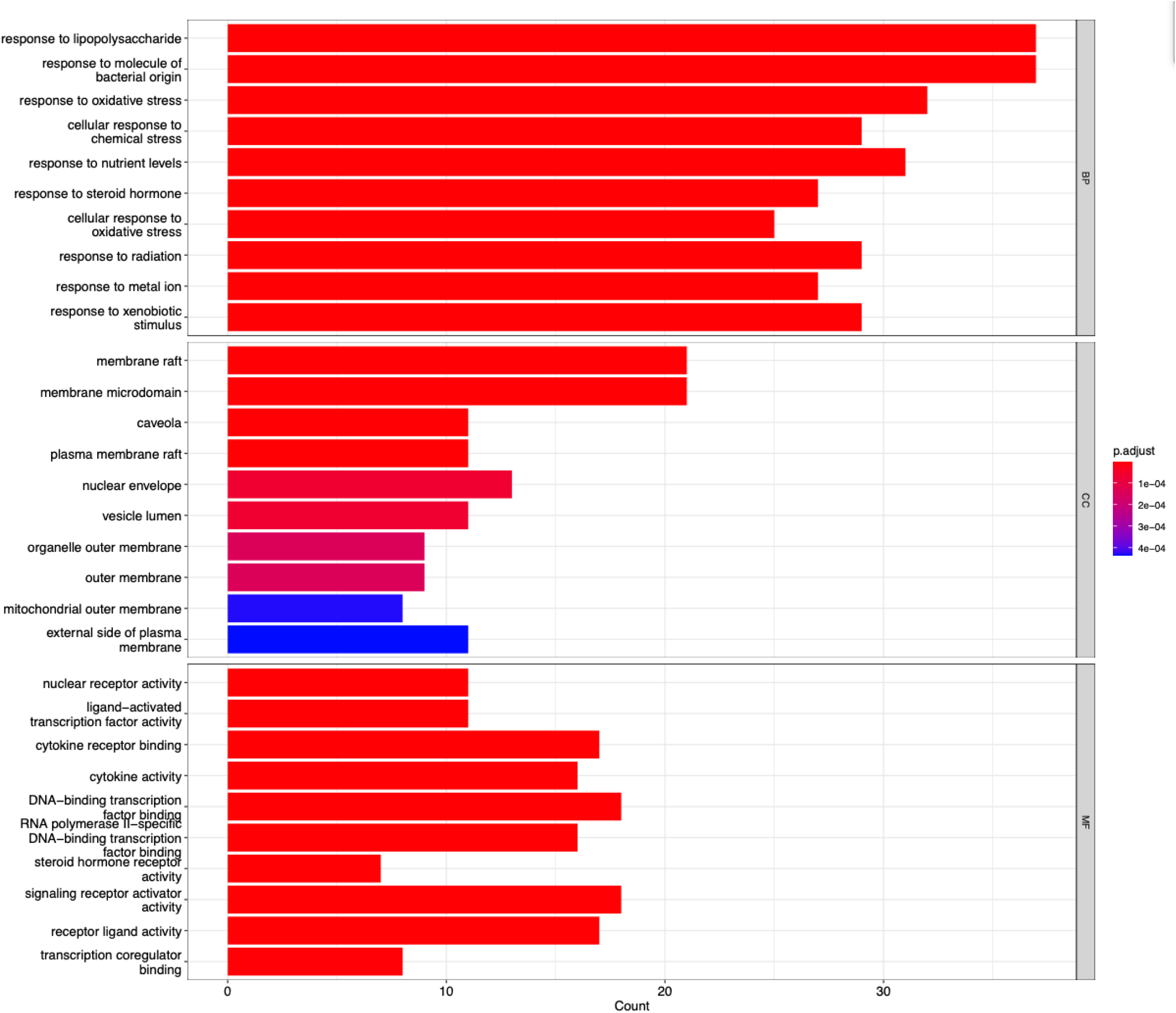
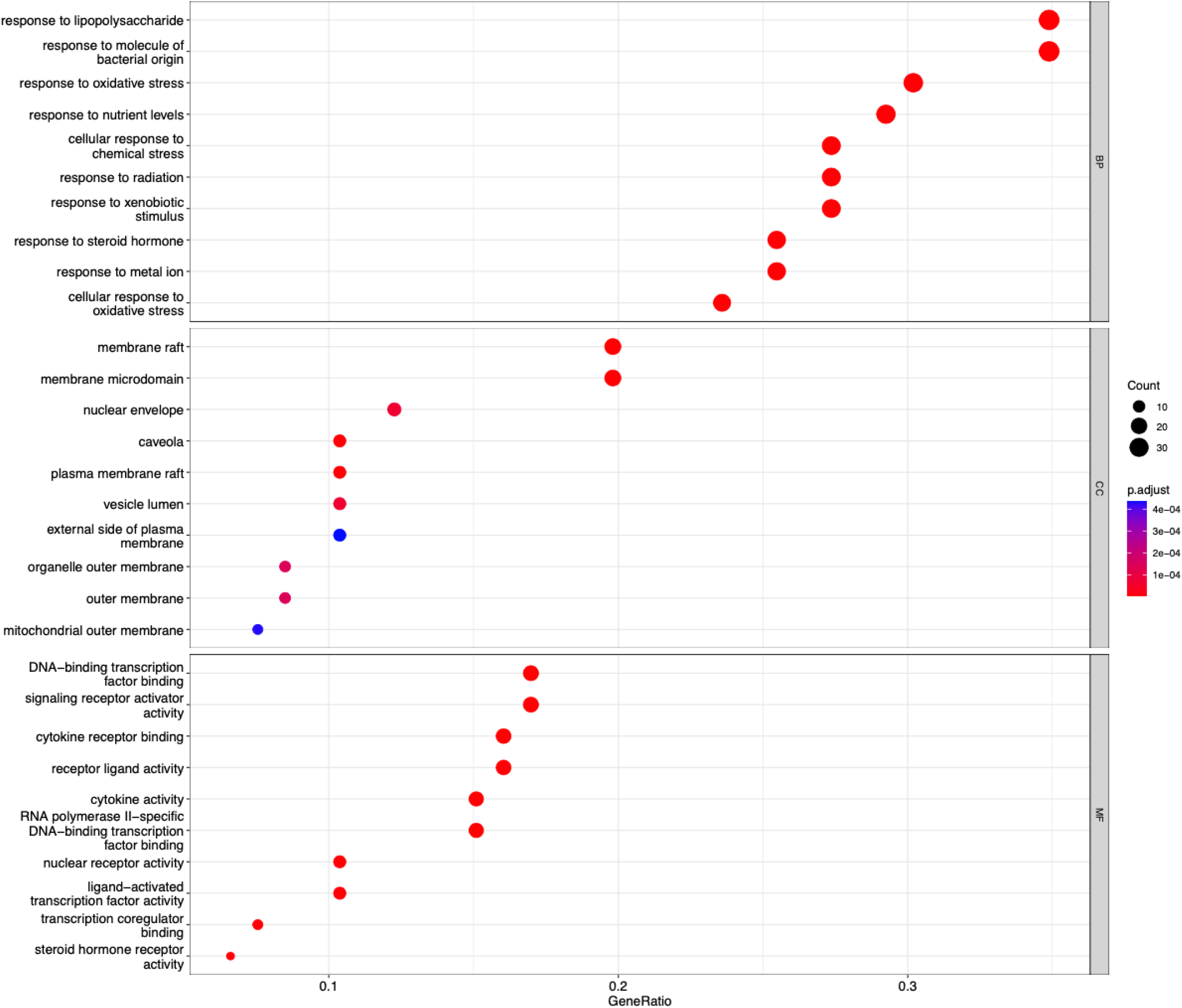
A. Barplot for Go enrichment analysis of DBG for SPID B. Bubble chart for Go enrichment analysis of DBG for SPID

#### KEGG Analysis

KEGG signaling pathway enrichment analysis showed that DBG treatment of SPID had a total of 152 signaling pathways, and the top 20 pathways were screened according to P<0.05, and the R language was used to draw a visual diagram of the target’s role in key pathways, as shown in Figs 6A and 6B. The main signaling pathways were PI3K-Akt, AGE– RAGE, IL-17, tumour necrosis factor (TNF), HIF-1 and Toll-like receptors signaling pathways, indicating that these key pathways can be used as the direction for the study of DBG in the treatment of SPID. As an example, the regulatory effects of these potential targets on the Toll-like receptors signaling pathway are presented in Figs 7A and 7B.

**Fig 6.**
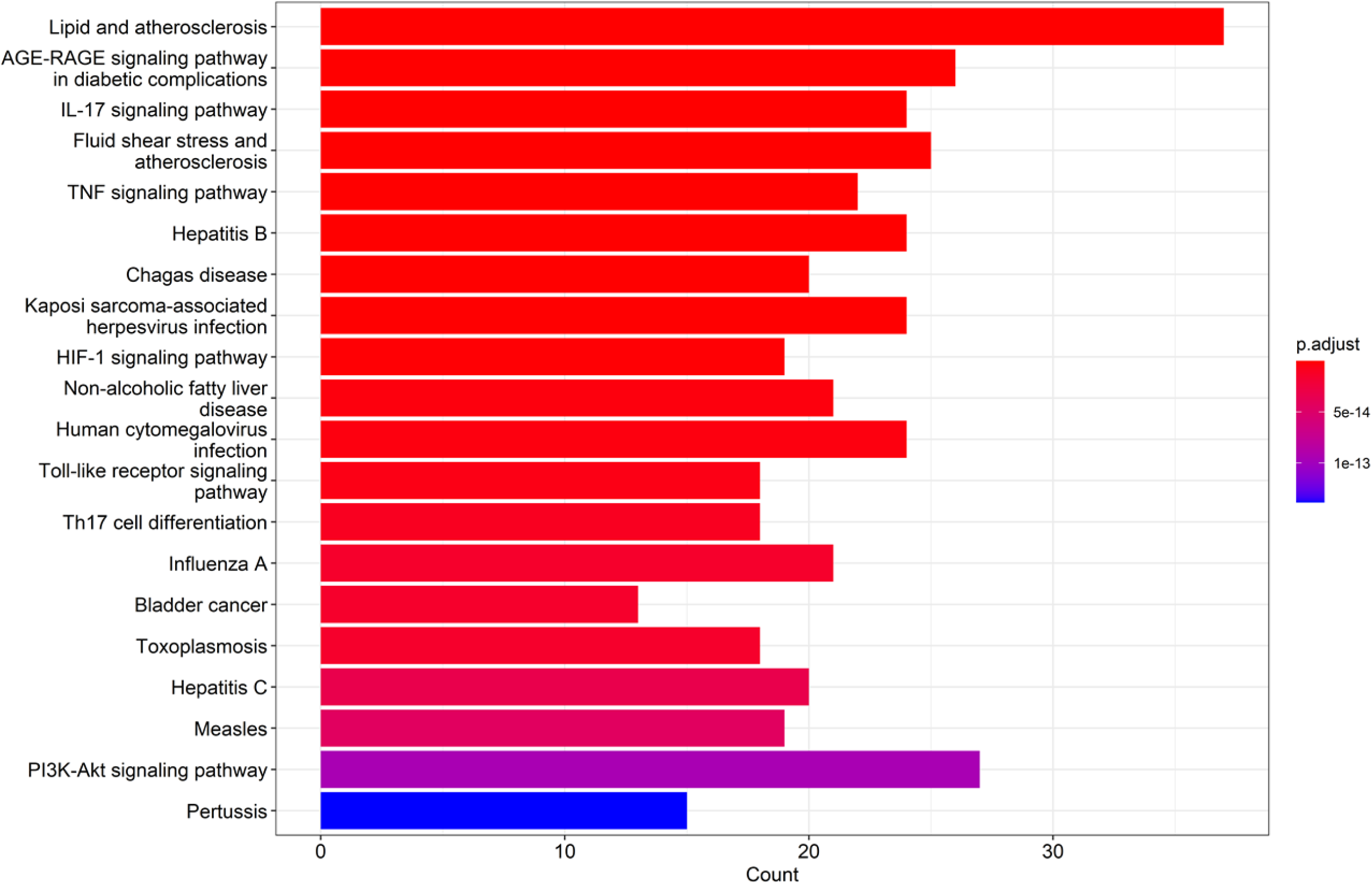
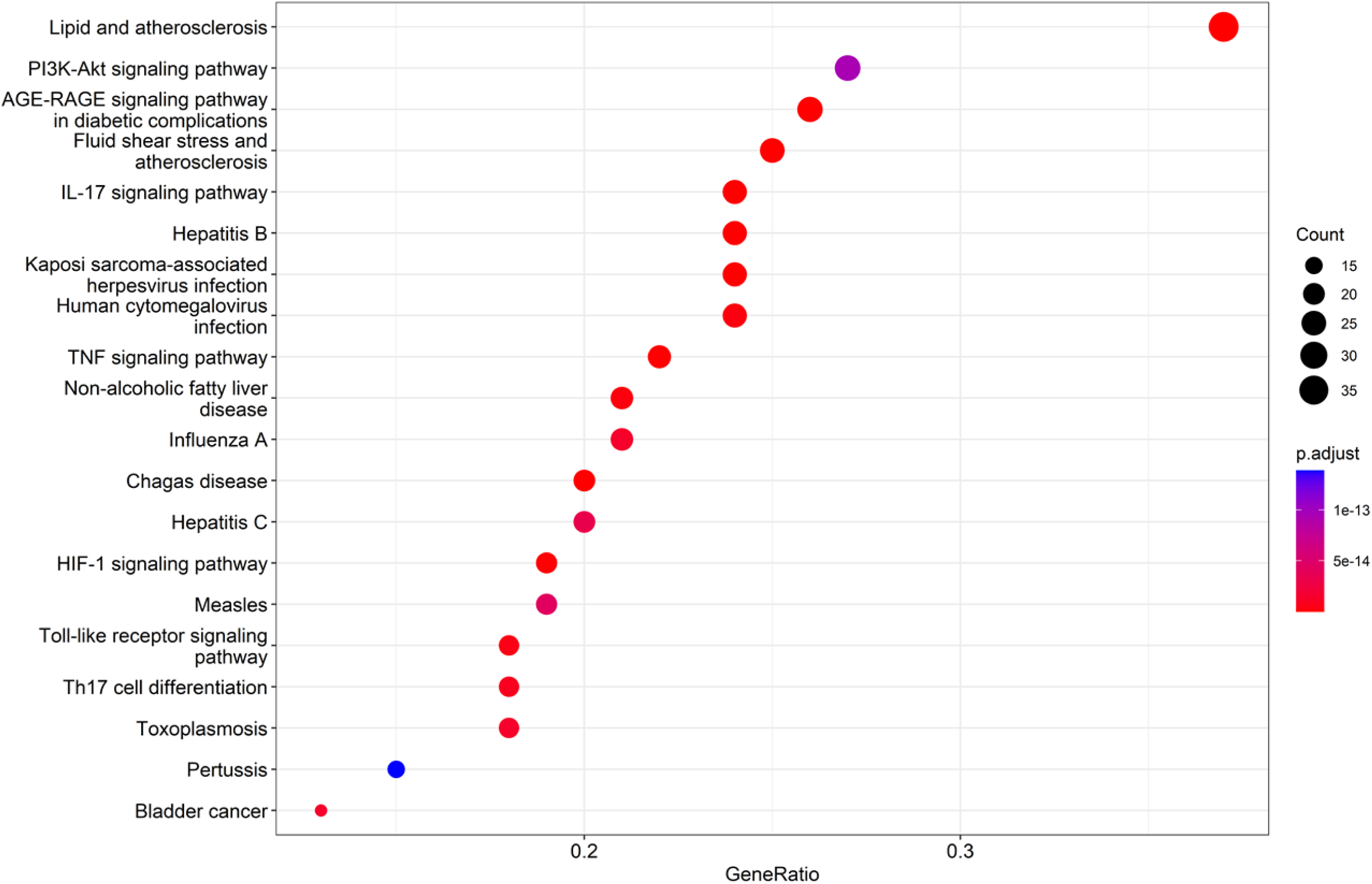
A. Barplot for KEGG enrichment analysis of DBG for SPID B. Bubble chart for KEGG enrichment analysis of DBG for SPID (top 20)

**Fig 7.**
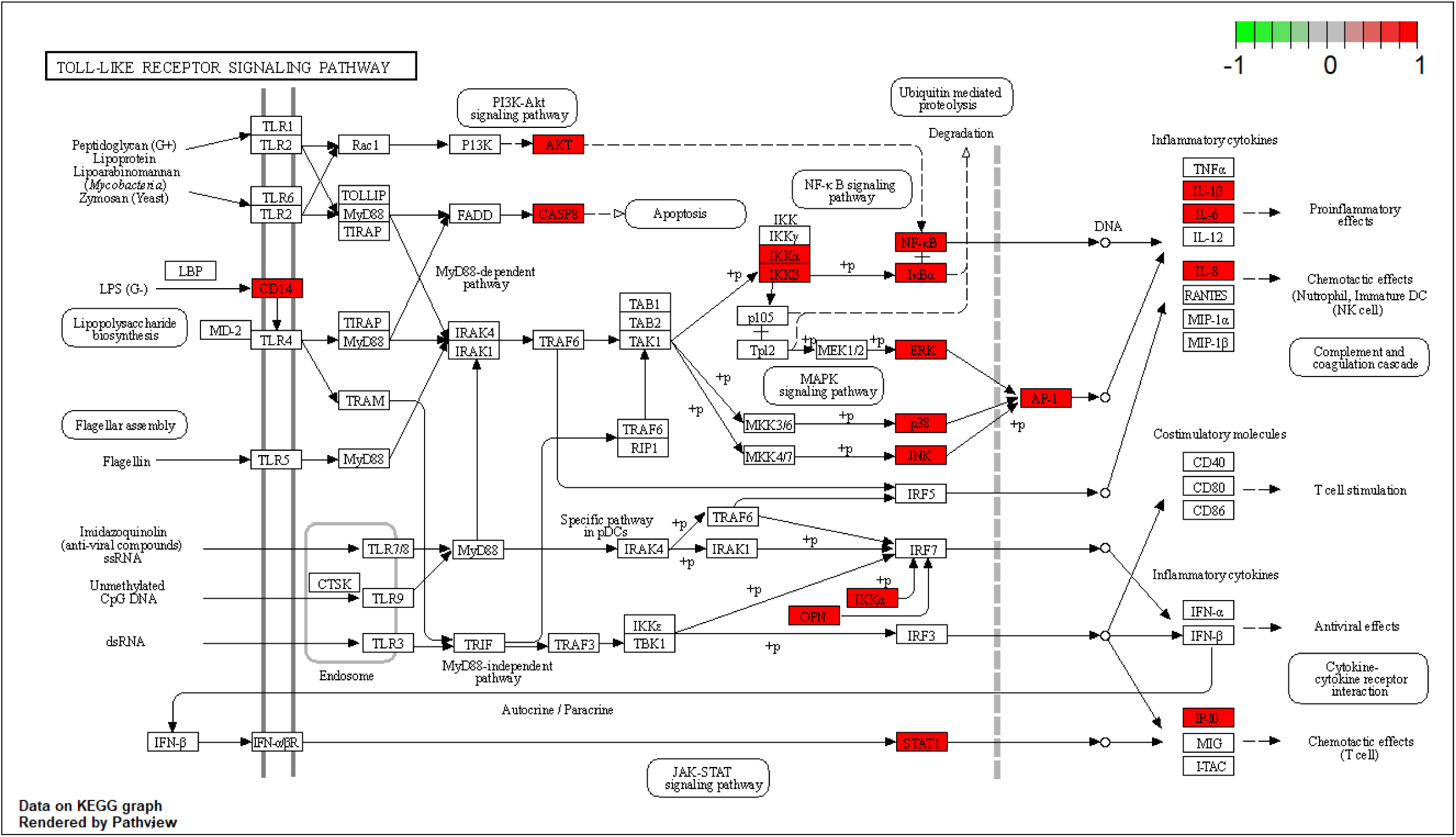
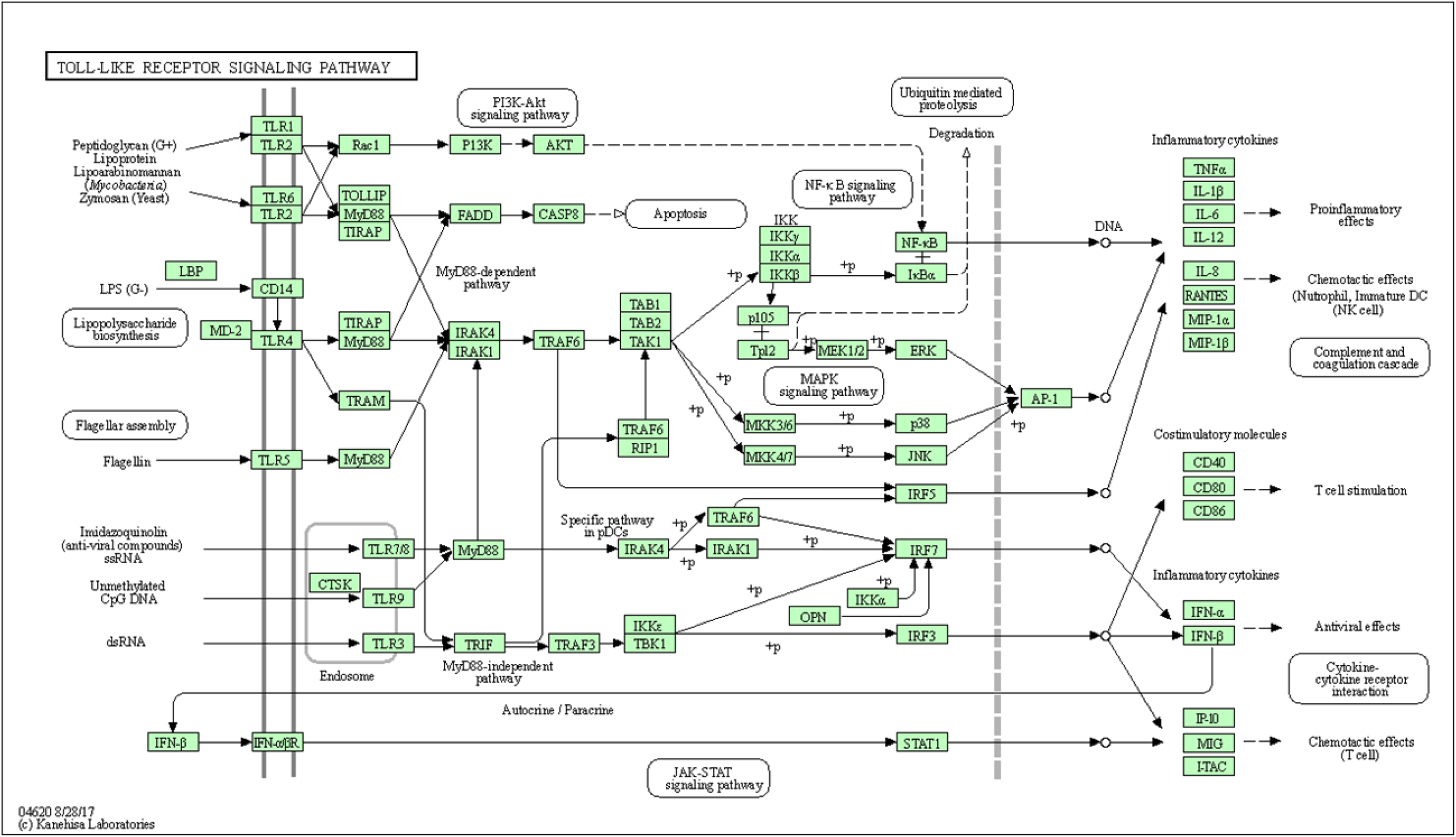
A. KEGG pathway analysis of Toll-like receptor signaling pathway of the DBG in the regulation of SPID B. KEGG pathway analysis of Toll-like receptor signaling pathway of the DBG in the regulation of SPID

### Results of Molecular Docking

Molecular docking was conducted for the top ten pharmacodynamic substances and the top six core genes screened by the PPI network with Autodock Tools 1.5.6. The protein IDs corresponding to the genes in the UniProt database and the corresponding proteins in the PDB protein database are shown in Table 2. The binding energy calculated by molecular docking are shown in Table 3. The lower the banding energy between ingredient and target, the better the combination effect between them. According to the above results, 15 pairs of target-ingredient combinations docking score are all lower than −5 kcal/mol, which indicates that they have stable binding properties. Active pockets were visualized using Pymol 2.3.1 software.

**Table 2:** Information of six proteins involved in molecular docking

| Targets | Uniprot-ID | PDB-ID | Ligand-ID |
| --- | --- | --- | --- |
| IL-6 | P05231 | 1ALU | SO4, LA |
| IL-1β | P01584 | 5R85 | S7A, SO4 |
| VEGFA | P15692 | 3QTK | ACT, GOL, TFA |
| CASP3 | P42574 | 1NME | 158, 159 |
| PTGS2 | P35354 | 5F19 | AKR, BOG, COH, EDO,<br>NAG |
| JUN | P05412 | 5FV8 | JEF |

**Table 3:** The binding energy of DBG core active ingredients and core common targets

| Core active ingredient | Banding energy(kcal/mol) |  |  |  |  |  |
| --- | --- | --- | --- | --- | --- | --- |
| | IL-6 | IL-1β | VEGFA | CASP3 | PTGS2 | JUN |
| Quercetin | -3.34 | -4.9 | -2.46 | -2.66 | -1.55 | -3.87 |
| Luteolin | -4.34 | -5.38 | -3.47 | -3.8 | -6.48 | -2.99 |
| Kaempferol | -4.26 | -5.08 | -2.7 | -2.71 | -2.5 | -2.65 |
| Beta-sitosterol | -4.31 | -6.56 | -2.88 | -3.55 | -2.4 | -3.24 |
| Naringenin | -3.35 | -5.41 | -3.7 | -2.53 | -2.71 | -3.21 |
| Stigmasterol | -5.08 | -7.21 | -3.68 | -4.18 | -2.74 | -4.51 |
| Acacetin | -3.57 | -5.52 | -2.9 | -3.36 | -7.14 | -3.59 |
| Formononetin | -3.91 | -5.76 | -3.52 | -3.18 | -4.13 | -3.83 |
| Diosgenin | -5.6 | -7.65 | -5.33 | -5.48 | -5.25 | -5.73 |
| Myricanone | -4.05 | -5.9 | -3.83 | -3.24 | -2.26 | -3.34 |

Detailed verification target-ingredient interactions were given in Fig 8.

**Fig 8.**
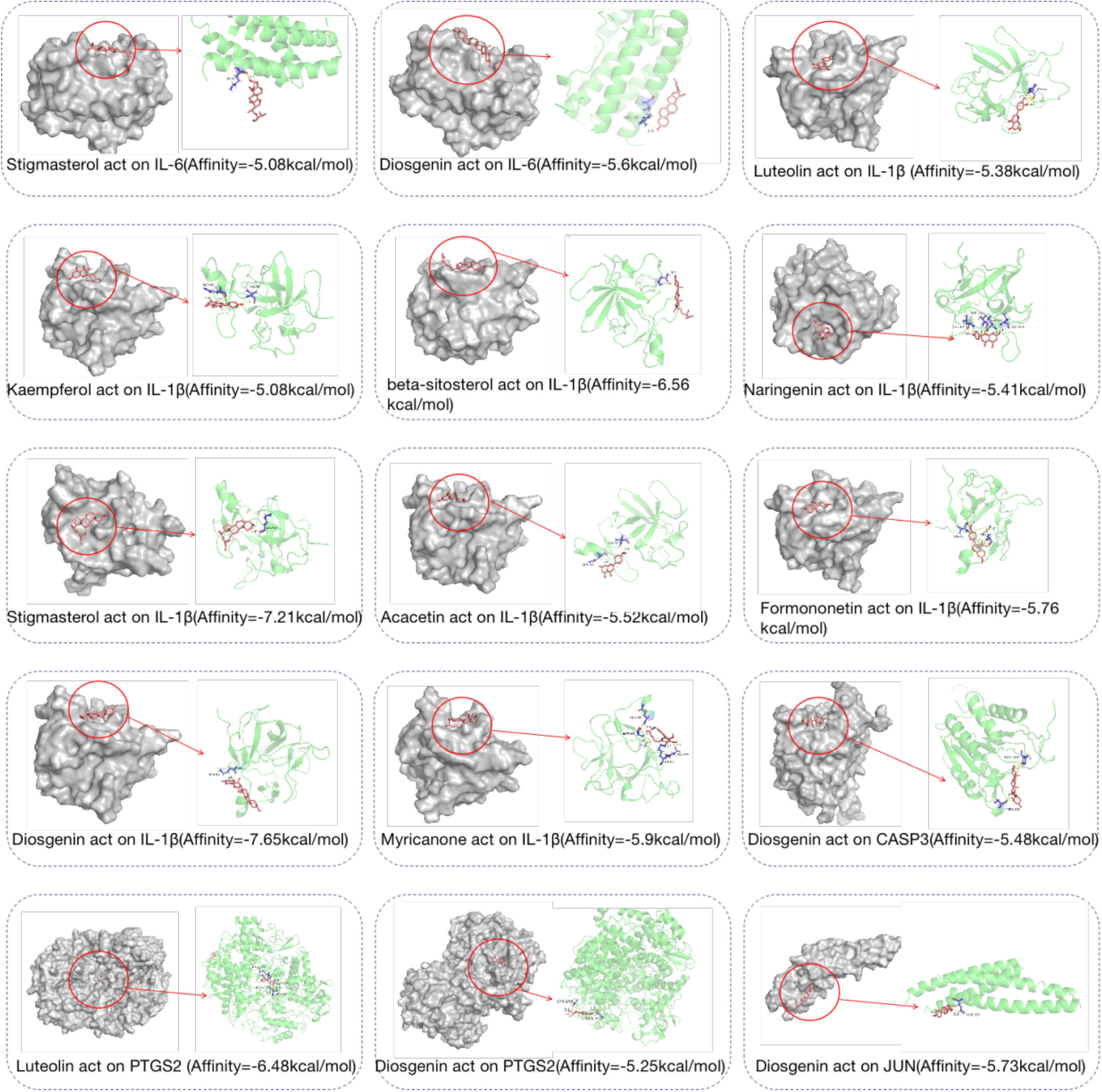
Detailed target-compound interactions of the molecular docking verification

### Effects on Inflammatory Factors

The changes of inflammatory factors in the serum of rats were detected by ELISA. Compared with the normal control group, the expressions of IL-6 and IL-1β in the model group were significantly increased (p<0.05), and the expressions of IL-4 and IL-10 in the model group were significantly decreased (p<0.05). indicating that there was an inflammatory response in the model group. Compared with the model group, the serum levels of IL-6 and IL-1β were significantly decreased (p<0.05) in each treatment group, while IL-4 and IL-10 were significantly increased (p<0.05), as shown in Table 4 and Fig 9.

**Fig 9.**
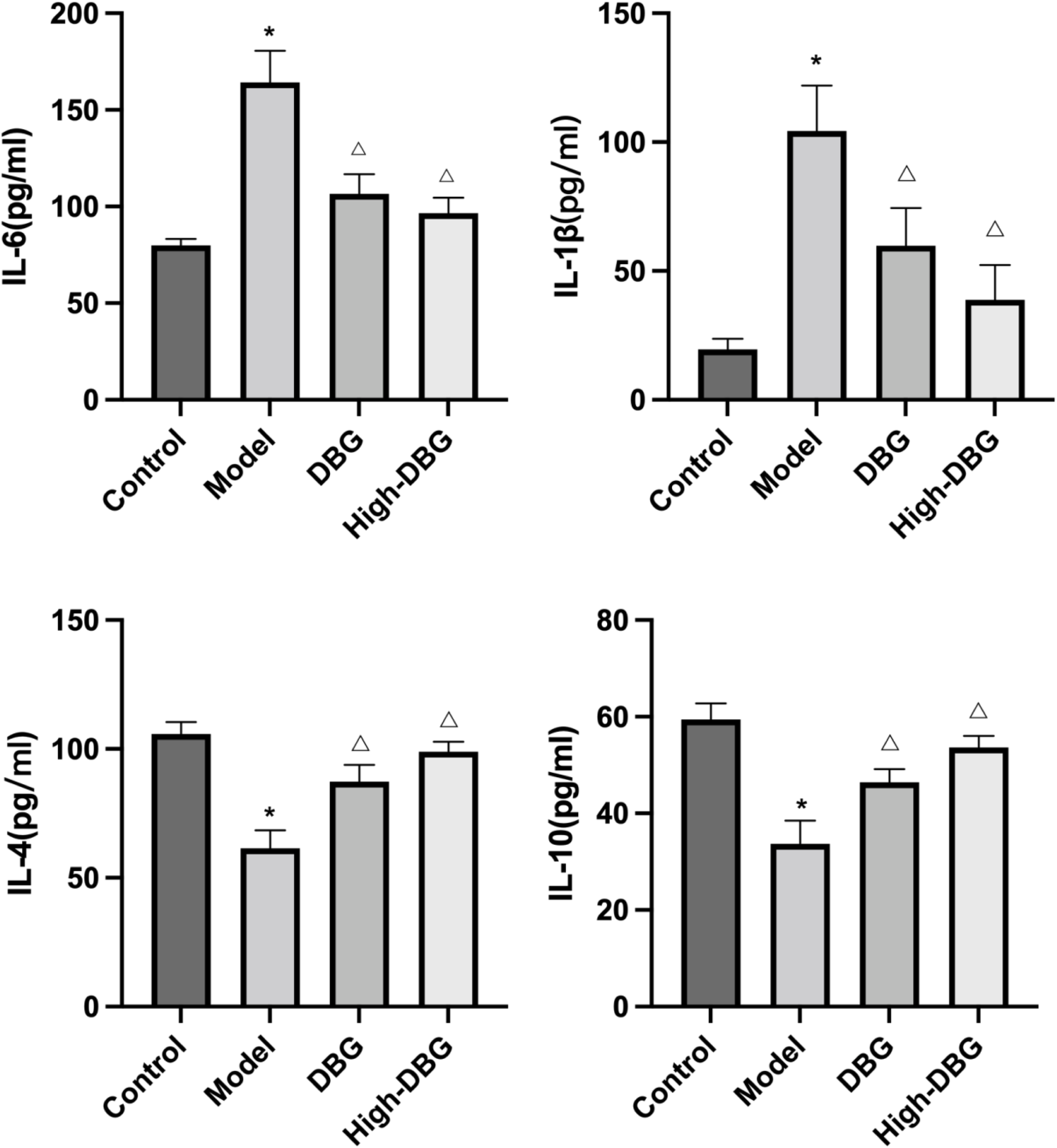
The serumIL-6, IL-1β, IL-4, IL-10 levels in SPID rats

**Table 4:** The serum IL-6, IL-1β, IL-4 and IL-10 levels in SPID Rats (Mean ± SD)

| | n | IL-6<br>(pg/mL) | IL-1β<br>(pg/mL) | IL-4<br>(pg/mL) | IL-10<br>(pg/mL) |
| --- | --- | --- | --- | --- | --- |
| control group | 9 | 79.93±3.33 | 19.61±1.68 | 105.84±4.55 | 59.46±3.03 |
| model group | 9 | 164.18±16.35* | 104.35±7.18 | 61.46±7.03* | 33.70±4.81* |
| DBG group | 9 | 106.61±10.13$^{\Delta}$ | 59.82±5.96 | 87.30±6.47$^{\Delta}$ | 46.44±2.74$^{\Delta}$ |
| High-DBG<br>group | 9 | 96.62±7.92<sup>Δ</sup> | 38.76±5.55 | 98.91±3.91<sup>Δ</sup> | 53.64±2.41<sup>Δ</sup> |

Notes: \*p<0.05 versus the model group; <sup>Δ</sup>p< 0.05 versus the model group.

### Western Blotting

Western blotting was performed to validate the TLR-4/MyD88/NF-κB p65 signaling pathway inferred from the KEEG results. As shown in Fig 10, compared with the control group, TLR-4, MyD88 and NF-κB p65 were significantly increased in the model group (p<0.05). Compared with the model group, the levels of TLR-4, MyD88 and NF-κB p65 were obviously downregulated in the DBG and High-DBG group (p<0.05). The above results revealed that DBG may treat SPID by modulating the TLR-4/MyD88/NF-κB p65 pathway.

**Fig 10.**
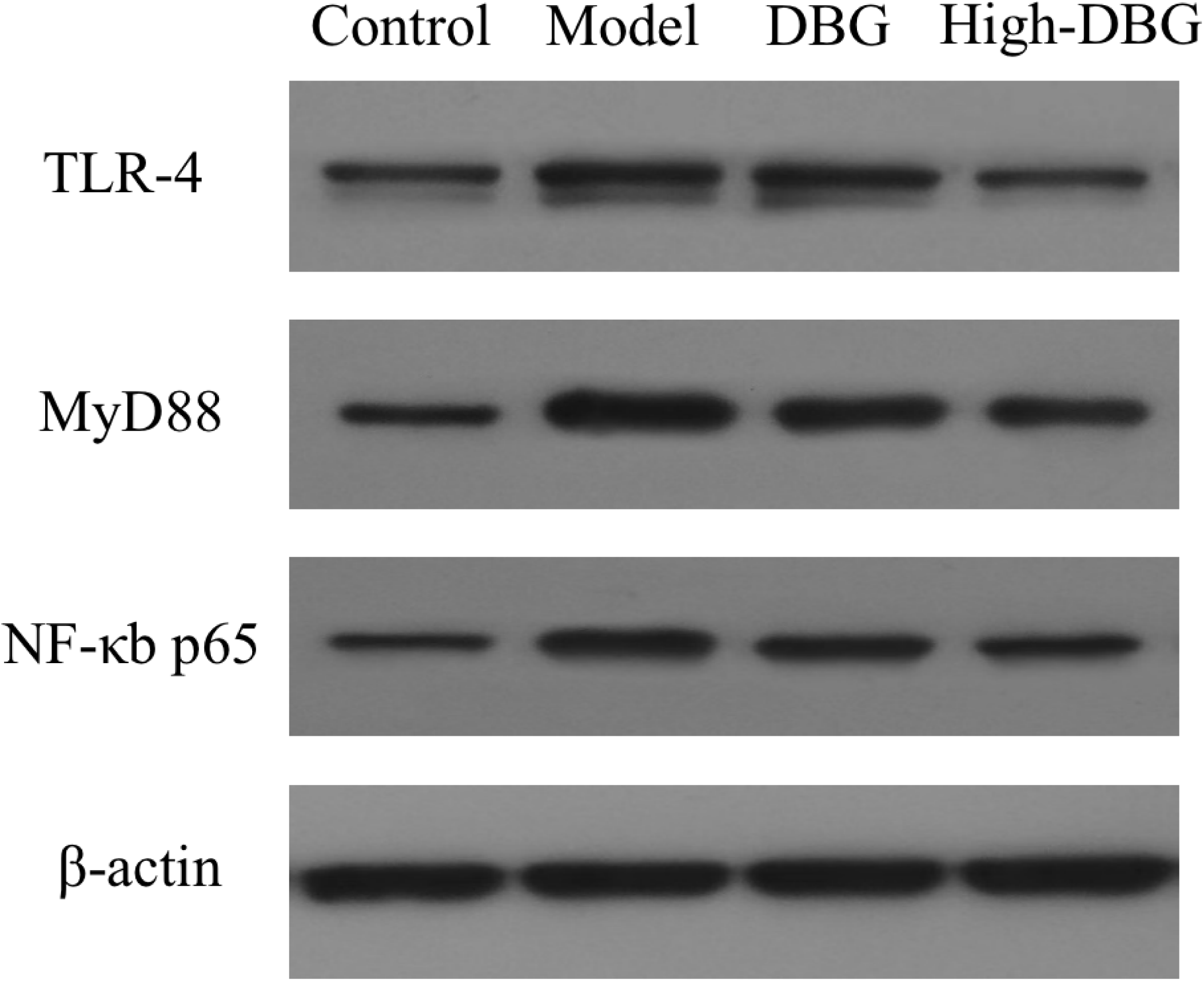
Effects of DBG on TLR-4/MyD88/NF-κB p65 signaling pathway related proteins in SPID rats

## Discussion

In this study, the potential mechanism of Danbai granules on pelvic inflammatory sequelae was clarified through network pharmacology and molecular docking methods combined with experimental verification. We screened out 46 active components of DBG, among which quercetin, luteolin, kaempferol, beta-sitosterol and naringenin were the most important chemical components. Through PPI network topology analysis and enrichment analysis, we found that the effective components in DBG may act on core targets such as IL-6, IL-1β, VEGFA, and CASP3 by regulating PI3K/AKT signaling pathway, AGE/RAGE signaling pathway, IL-17 signaling pathway, TNF-α signaling pathway, HIF-1 signaling pathway, Toll-like receptor signaling pathway, etc. to effectively treat SPID. In addition, through experimental evidence, we confirmed that DBG may inhibit pelvic inflammatory response by regulating Toll-like receptor 4/MyD88/NF-κb p65 signaling pathway, so as to achieve the purpose of treating SPID. These results highlight the multi-component, multi-target and overall regulation characteristics of traditional Chinese medicine preparations, and further provide theoretical support for the treatment of SPID.

We screened out the core compounds of Danbai Granules through R language, including quercetin, luteolin, kaempferol, β-sitosterol, etc., which were likely to be the main components of DBG in the treatment of SPID. The anti-inflammatory mechanism of quercetin is complex. It has significant anti-inflammatory effects in both acute and chronic inflammatory conditions [35]. A study has shown that quercetin has strong anti-inflammatory and antioxidant effects by ERK1//2 MAPK and NF-κb pathway through regulating the expression of multiple inflammatory cytokines [36]. Jingwen Zhang observed that quercetin can exhibite anti-inflammatory effects by reducing the expression levels of TNF-α, IL-6 and IL-1β in LPS-induced RAW264.7 macrophages and regulate the PI3K/Akt signaling pathway [37]. Luteolin belongs to the flavone family commonly present in plants. Studies have shown that both isolated luteolin and plant extracts have anti-inflammatory activity [38–39]. Luteolin-7-O-Glucuronide (L7Gn), an active component of luteolin, has been identified to prevent overproduction of inflammatory mediators including IL-6, IL-1β and cyclooxygenase 2 [40]. JunKyung Lee has found that luteolin inhibited the expression levels of TLR3 or TLR4 target genes, including TNF-α, IL-6, IL-12 and IL-27 [41]. and that luteolin could effectively eliminate the inflammatory process activated by keratitis by inhibiting TLR3/TAK/NF-κB signaling pathway [42]. Kaempferol is well known for the desirable properties in radical scavenging, oxidation resistance and inflammation resistance. It can partly adjust the immune response through T-cell response [43]. Kaempferol derivatives can be used in clinic as antimicrobial agents [44]. Yifei Bian has found that kaempferol can significantly reduce LPS-induced tumor necrosis factor-α (TNF-α), interleukin-1 (IL-1), interleukin-6 (IL-6), Overproduction of intercellular adhesion molecule-1 (ICAM-1) and vascular cell adhesion molecule-1 (VCAM-1), and kaempferol is negative for Toll-like receptor 4, NF-κB and STAT signaling Regulation [45]. Anti-inflammatory, immunomodulatory and analgesic effects of beta-sitosterol have also been identified [46]. β-Sitosterol can inhibit inflammatory factors such as TNF-α, IL-1β, IL-6, IL-8 and ROS, reduce the expression of key components of the NLRP3 inflammasome and inhibit the activation of caspase-1 [47].

This study found that IL-6, IL-1β, VEGFA, CASP3, PTGS2, JUN, etc., are potential targets of Danbai Granules in the treatment of SPID by constructing a PPI network of active targets. Both IL-6 and IL-1β play a role in promoting inflammatory response and aggravating inflammatory infiltration of tissues [48]. IL-6 is a cytokine in the chemokine family, which acts on numerous target cells, including macrophages, stationary T cells, activated B cells and plasma cells, to induce the migration of these immune cells to the lesionto phagocytose or kill pathogens [49–50], IL-6 can combine with neutrophils to change the morphology of neutrophils, thereby activating the inflammatory response of the body and intensifying histopathological damage [51]. The level of IL-1β in normal tissues is very low, when injury or inflammation occurs, IL-1β will be produced in large quantities and further induces the release of downstream proinflammatory factors to participate in the occurrence and expansion of SPID inflammation [52]. Vascular endothelial growth factor VEGF is the strongest angiogenesis stimulator, and its main role in endometrium is to promote the proliferation of vascular endothelial cells and and increase vascular permeability, Its abnormal expression is involved in intrauterine adhesions, endometrium. fibrosis and other pathological processes [53]. The process of inhibition and elimination of inflammation is closely related to apoptosis. When an inflammatory response occur, CASP3 can activate the Bcl-2 family and to induce apoptosis of pro-inflammatory cells, thereby inhibiting the inflammatory response and facilitating the absorption and dissipation of inflammation [54]. Studies have shown that caspase-3 and Caspase-8 are involved in the regression of chronic pelvic inflammatory disease in rats [55].

In this study, GO enrichment and KEGG pathway enrichment analysis were performed to elucidate various mechanisms of treatment of SPID by DBG. The bubble chart showed that DBG mainly treats SPID through biological processes. BP entries include response to lipopolysaccharide, response to molecule of bacterial origin, and response to oxidative stress, which further play a pharmacological role in inhibiting inflammation and relieving pain. Furthermore, these putative target genes were selected for KEGG pathway analysis, and a total of 20 important pathways that may be controlled by DBG in the treatment of SPID were discovered. The results showed that the main signaling pathways involved in DBG treatment of SPID included cancer signaling pathways, infectious disease-related signaling pathways, apoptosis, PI3K-Akt, Toll-like receptors, etc. These KEGG signaling pathways are not entirely independent but are generally related. They play a synergistic role in the pathogenesis and treatment of SPID, and also provide a strong basis for TCM multi-target therapy. There are pathways related to body immunity. For instance, Toll-like receptors (TLRs) can promote the initiation and occurrence of the body’s innate immune response, and at the same time can participate in the activation of partial acquired immunity, which are known to play a role in chronic pain [56]. Activation of TLR4 recruits myeloid differentiation protein 88 (MyD88), an intracytoplasmic downstream adaptor molecule, which activates inflammation, proliferation and apoptosis. A Study have shown that Among African American women with PID, variants in the TLR1 and TLR4 genes, which may increase signaling, were associated with increased C. trachomatis infection [57]. Sex steroids may have complex effects on TLR4 signaling, causing chronic inflammation and chronic pain in women [58].

Molecular docking calculation results show that some components in DBG can be docked with IL-6, IL-1β, VEGFA, CASP3, PTGS2, JUN, and the binding energy of Stigmasterol to IL-1β, Diosgenin to IL-1β are ≤ -7.0 kcal/mol, indicating that these interactions is very important. It may be the main component of the interaction between DBG and SPID. Prompted by the predicted results of network pharmacology and molecular docking. Prompted by the prediction results of network pharmacology and molecular docking, this study was verified by an animal experiment. The experimental results showed that compared with the control group, the contents of IL-6 and IL-1β were drastically increased (P<0.05) in the model group, while the expressions of IL-4 and IL-10 was significantly decreased (P<0.05), and the protein expressions of TLR4, MyD88 and NF-κB p65 were significantly increased (P<0.05). Compared with the model group, the serum levels of IL-6 and IL-1β in each treatment group were significantly decreased (P<0.05), while the levels of IL-4 and IL-10 were significantly increased (P<0.05). TLR4, MyD88, and NF-κB p65 protein expressions were significantly decreased (P<0.05), suggesting that DBG may down-regulate inflammatory factors in serum of rats with SPID and inhibit the expression of TLR4/MyD88/NF-κB p65 signaling pathway-related proteins expression, reduce pelvic inflammatory response to reat the SPID effectively. This is also consistent with the predicted results of network pharmacology. Hence, DBG plays a vital role in the treatment of SPID through a multi-target and multi-channel network approach, and it is necessary to further explore the molecular mechanism of DBG in the treatment of SPID in vitro and in vivo.

## Conclusion

Based on the network pharmacology approach, we have screened out quercetin, luteolin, kaempferol, beta-sitosterol, naringenin, stigmasterol, acacetin, formononetin, diosgenin, myricanone, and other potentially active ingredients. At the same time, the hub genes IL-6, IL-1β, VEGFA, CASP3, PTGS2, JUN, etc. were screened. Go and KEGG enrichment analysis showed that Toll-like receptor signaling pathway plays an important role in DBG regulation of SPID, which was verified in vivo experiments. The application of molecular docking technology enables the binding activity of active ingredients to receptor proteins to be confirmed. providing a theoretical support and scientific basis for the clinical application of DBG to treat SPID. This study provides new enlightenment for the application of Danbai granules and the treatment of SPID.

## Data Availability

The data used to support the findings of this study are available from the corresponding author upon request.

